# The inferior olive transforms upstream sensorimotor errors into cerebellar teaching signals

**DOI:** 10.64898/2026.03.11.711185

**Authors:** Pierce Mullen, Hesho Shaweis, Maarten Zwart

## Abstract

Adaptive behavior requires evaluating whether sensory feedback matches expectations derived from motor commands. In cerebellar theories, climbing fibers arising from the inferior olive (IO) convey prediction error signals that instruct learning, yet how these signals are constructed remains unresolved. Using larval zebrafish performing visuomotor behaviors in virtual reality, we combined two-photon calcium imaging with glutamate sensors to measure excitatory input to the IO and climbing fiber output under identical conditions. IO activity scaled with discrepancies between expected and experienced optic flow across open- and closed-loop perturbations. Strikingly, excitatory drive to the IO already contained structured, motor-dependent information consistent with error-like signals. Comparison of input and output revealed selective, frequency-dependent transmission from the olive to the cerebellum. These findings suggest where error representations arise in the olivo-cerebellar pathway and support a model in which climbing fiber teaching signals reflect upstream processing combined with filtering within the IO.

## Introduction

To adapt to dynamic environments, animals must learn from the sensory consequences of their actions. Such learning requires distinguishing self-generated sensory feedback from externally generated stimuli and determining when action outcomes deviate from internal expectations. These deviations are often formalized as prediction errors, signals reflecting mismatches between expected and experienced feedback that can instruct adaptive changes in future behavior. In motor control, prediction error-like computations are central to theories of cerebellar learning and internal model updating ^1–3^ .

A major behavioral adjustment occurs when animals modify motor vigor or timing in response to changes in action effectiveness. Larval zebrafish, for example, adapt swimming to stabilize position under altered visual feedback gain or temporal delays in reafference ^4–6^. Such learning requires monitoring the sensory outcomes of locomotion while filtering sensory fluctuations unrelated to action. More generally, computing action outcomes depends on integrating motor commands with sensory feedback to infer which sensory events are explainable consequences of behavior and which represent unexpected perturbations that should drive learning.

The cerebellum has long been proposed to support these computations through experience-dependent refinement of sensorimotor transformations. Classical models posit that climbing fibers, originating from the inferior olive (IO), provide an instructive teaching signal that guides plasticity in cerebellar cortex ^1–3^. Because climbing fiber activity is modulated by unexpected sensory events and perturbations, these signals are widely interpreted as encoding errors or prediction error-like quantities ^7,8^. However, a critical gap remains: how are these instructive signals constructed within the olivo-cerebellar pathway?

The IO has been postulated to compute discrepancies between intended and actual outcomes and relay prediction errors to the cerebellum ^1–3^. At the same time, the IO receives diverse excitatory afferents and strong inhibitory feedback from cerebellar nuclei that can gate olivary output and synchrony ^9^. Moreover, the relative timing of excitation and inhibition can shape the probability and timing of olivary spikes and thus climbing fiber transmission ^10,11^. Recent in vivo imaging studies have begun to reveal how IO neurons integrate excitatory and inhibitory inputs to generate olivocerebellar signals ^12^. These observations raise a fundamental question: does the IO generate error structure *de novo*, or does it receive excitatory drive already shaped by upstream sensorimotor processing and selectively transmit it to the cerebellum?

Testing this question directly has been difficult. Error-related signaling in the olivo-cerebellar system is most often inferred from Purkinje-cell complex spikes or CF-driven calcium signals, which provide high-resolution access to pathway output but not the excitatory synaptic drive impinging on IO neurons ^7,8,13,14^. Conversely, mechanistic insight into how excitation, inhibition, and electrical coupling gate IO spiking derives largely from reduced preparations in which inputs are electrically evoked or optogenetically controlled, limiting insight into the structure of behaviorally generated input patterns ^9,11,15,16^. The *in vivo* transformation from excitatory synaptic drive in the IO to climbing fiber teaching signals therefore remains largely unobserved.

Larval zebrafish provide the means to bridge this gap. The IO and cerebellum are optically accessible at cellular resolution, and sensorimotor behaviors such as the optomotor response can be studied in closed-loop virtual environments where visual feedback can be precisely perturbed ^4,5^. Recent work has established the structural and functional organization of visually tuned responses in the zebrafish IO, providing a foundation for probing how error-like signals emerge within this circuit ^17^.

In this study, we investigate how prediction error-like signals, here defined as discrepancies between expected and experienced visual feedback on behavior, are represented and transformed within the IO during visuomotor learning in larval zebrafish. Using two-photon calcium imaging during open-loop and closed-loop perturbations, we find that IO activity reflects the sign and amplitude of the discrepancy between expected and actual visual feedback across contexts. We then directly measure excitatory synaptic drive to the IO using iGluSnFR ^18,19^ in vGluT2-positive olivary neurons and compare it to glutamate release from their climbing fiber terminals under identical behavioral conditions. This input-output comparison reveals structured, motor-dependent information in excitatory drive to the IO and shows that climbing fiber output reflects selective, frequency-dependent filtering of that drive across visuo-motor space and timescales. Together, our results help define the locus of error construction in the olivo-cerebellar system and support a model in which climbing fiber teaching signals emerge through a combination of upstream shaping and olivary filtering rather than computation solely within the IO.

## Results

### IO neurons encode prediction errors as sensory discrepancies

We first asked whether neurons in the IO do encode the discrepancy between expected and actual outcomes (Fig 1B). To understand what a prediction error looks like to a zebrafish, we analyzed a behavioral dataset ^20,21^ containing freely swimming zebrafish larvae and plotted the relationship between visual scene changes and proxies of swim power such as tail beat frequency (Fig 1A). This revealed an inverse sigmoid-like curve where the fish expects an increasingly negative velocity of the visual field (backwards motion) with stronger swims. This curve, representing the ethological sensory expectation for a given motor swim command, could then be used to define a series of hypotheses that error-encoding neurons should satisfy when actual sensory experiences deviated from the expectation curve. The first prediction of this hypothesis was that a neuron encoding prediction errors should report the absolute discrepancy between the expected reafferent velocity, and the actual velocity experienced by the fish.

**Figure 1.**
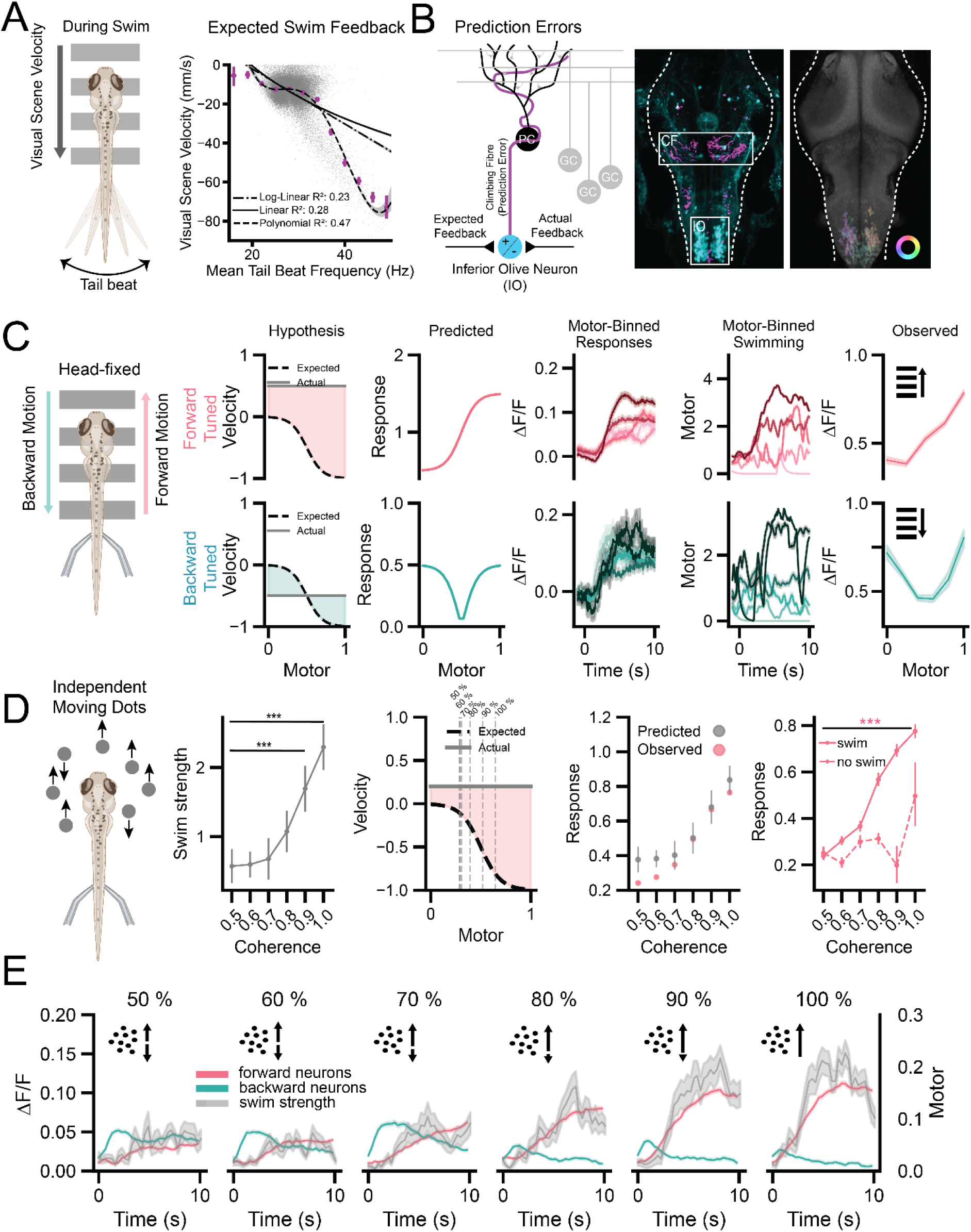
IO responses in the absence of reafferent feedback. A) Left panel: freely swimming zebrafish larvae experience different degrees of movement of the visual scene for different swim strengths. Right panel: The relationship between mean tail beat frequency (Hz) and velocity of the visual scene during forward OMR-induced swims in freely swimming zebrafish larvae. B) Left panel: the comparator hypothesis – inferior olive neurons contrast expected visual feedback with actual visual feedback to compute and send prediction errors to the cerebellum. Middle panel: A maximum projection of a zebrafish whole brain volume from a transgenic line expressing Kaede (y330:UAS:kaede) showing IO nuclei location in zebrafish. Cell bodies in the IO (cyan) and climbing fibers projecting to the cerebellum(magenta) have been differentially color-coded for clarity. Right panel: Spatial organization of directional tuning in the IO. C) Left panel: Moving gratings were projected at a constant velocity beneath head-fixed, immobilized zebrafish larvae whilst recording fictive swims from the zebrafish tail. 2^nd^ panel: The hypothesis for forward-tuned error neurons (top) and backward-tuned error neurons (bottom) with an expectation curve during either a constant forward or backward velocity, respectively. 3^rd^ panel: The expected response proportional to the difference between the expectation curve and actual constant velocity. 4^th^ panel: Recorded direction-tuned IO neuron responses binned by swim power (5^th^ panel). Right panel: The observed relationship of recorded IO responses as a function of swim power at a constant velocity. D) Left panel: Moving dots were presented to zebrafish larvae with different proportions moving in the forwards versus backwards direction. 2^nd^ panel: Coherence of 1 (100%) corresponds to all dots moving in the forward direction. The relationship between moving dot coherence and fictive swim strength. 3^rd^ panel: Average swim strength for each coherence across trials was used to identify the expected velocity to contrast with the actual, constant, velocity. 4^th^ panel: The predicted and observed responses for forward-tuned neurons. Right panel: A comparison between forward-tuned responses at each coherence level during trials that did and did not have swims. E) Average trial responses of forward-tuned and backward tuned neurons at each coherence level. Swim strength (Motor) is in grey.

Consequently, when fish experience a constant velocity of visual flow in the absence of reafference (open-loop), IO neuron responses should vary only as a function of swim strength. We confirmed prior work showing that IO neurons exhibit direction-selective visual flow responses (Fig 1B ^17^) and simplified our investigation to forward-tuned and backward-tuned IO neurons in a 1-dimensional forwards and backwards visual motion behavioral paradigm. Accordingly, we predicted that IO neurons tuned to forwards visual flow should increase non-linearly with swim strength when visual flow is forward-moving, whereas IO neurons tuned to backward visual flow should exhibit a bi-directional relationship when visual flow is backward moving (Fig 1C, 2^nd^, 3^rd^ panel). To experimentally test this, we projected a series of moving visual gratings below the fish whilst recording IO calcium dynamics using two-photon microscopy and zebrafish expressing nuclear pan-neuronal GCaMP6f (HuC:H2B-GCaMP6f). In this open-loop paradigm, fish experience zero reafference, allowing us to measure IO activity as a function of swim strength in the absence of self-generated visual feedback. We calculated the direction-tuning of neurons and measured the swim power from electrophysiological recordings of fictive swims from the zebrafish tail (Fig 1C, 1^st^ panel). Binning trials by swim power and plotting against the IO calcium dynamics indeed revealed a positive non-linear relationship for forward-tuned IO neurons and an approximately u-shaped relationship for backward-tuned neurons, consistent with the hypothesis of error encoding (Fig 1C).

Zebrafish swim power is modulated by the certainty of visual stimuli ^22^, so to directly test the effect of swim strength on IO responses we introduced different degrees of uncertainty in the visual stimulus. We projected a visual scene of dots moving at the same velocity in the forwards direction and adjusted the proportion of dots moving in the forwards direction versus the backwards direction; all dots, or half the dots, moving in the forward direction corresponded to 100%, and 50%, stimulus coherence, respectively. This decoupled swim power from the velocity of the visual scene, where swim power significantly increased with stimulus coherence (mixed-effect linear model; coherence (1.0 vs 0.5): β = 1.664, 95% CI [1.1, 2.228], p < 0.001) (Fig 1D). We used swim strength values from each coherence trial to hypothesize the sensory discrepancy that error-encoding neurons should report during this experiment (Fig 1D, 3^rd^ panel). In line with this hypothesis, stimulus coherence modulated forward-tuned neuron activity during swimming, with higher activity at 100 % coherence than at 50 % coherence (mixed-effects linear model; coherence × swim interaction: β = 0.259, 95% CI [0.159, 0.360], p < 0.001) (Fig 1D, 4^th^, 5^th^ panel).

Conversely, as the proportion of dots moving in the backward direction increased, we saw a proportional rapid activation of backward-tuned neurons that became suppressed on the initiation of swimming and forward-tuned neuron activity (Fig 1E). Together, these initial findings using constant velocity in open-loop were consistent with IO neurons encoding sensorimotor prediction errors as sensory discrepancies.

### IO neurons report magnitude of perturbation in closed-loop feedback

We next confirmed whether our findings were consistent when the zebrafish could interact with the visual scene, in the context of both normal reafference and altered reafference or exafference. We performed closed-loop experiments where fictive swims recorded from the tail could drive the visual scene, providing reafference feedback signals to the zebrafish that we controlled through a feedback gain parameter ^4^ (Fig 2A). A series of ten repetitions of two-second long forward moving gratings were presented to the fish to establish a baseline, followed by three repetitions where either the closed-loop feedback gain was altered (reverse/no reafference, gain up or gain down) or the velocity of the visual scene was altered (+ve, -ve probe) (Fig 2B, C). We used the same sigmoidal velocity expectation as before and modelled the actual velocity as a combination of constant velocity (the stimulus) and a negative linear velocity proportional to swim strength (the reafference) to represent the linear closed-loop feedback relationship in the virtual reality setup (Fig 2A). This predicted that reversing the sign of reafference should produce the largest forward error and thus the largest response in forward-tuned neurons, and that backward-tuned neurons would still exhibit a u-shaped relationship with swim power during backward-moving visual scenes (Fig 2D). Indeed, we found a striking similarity between the model predictions and observed response magnitudes (Fig 2E) and relationship with swim power (Fig 2D).

**Figure 2.**
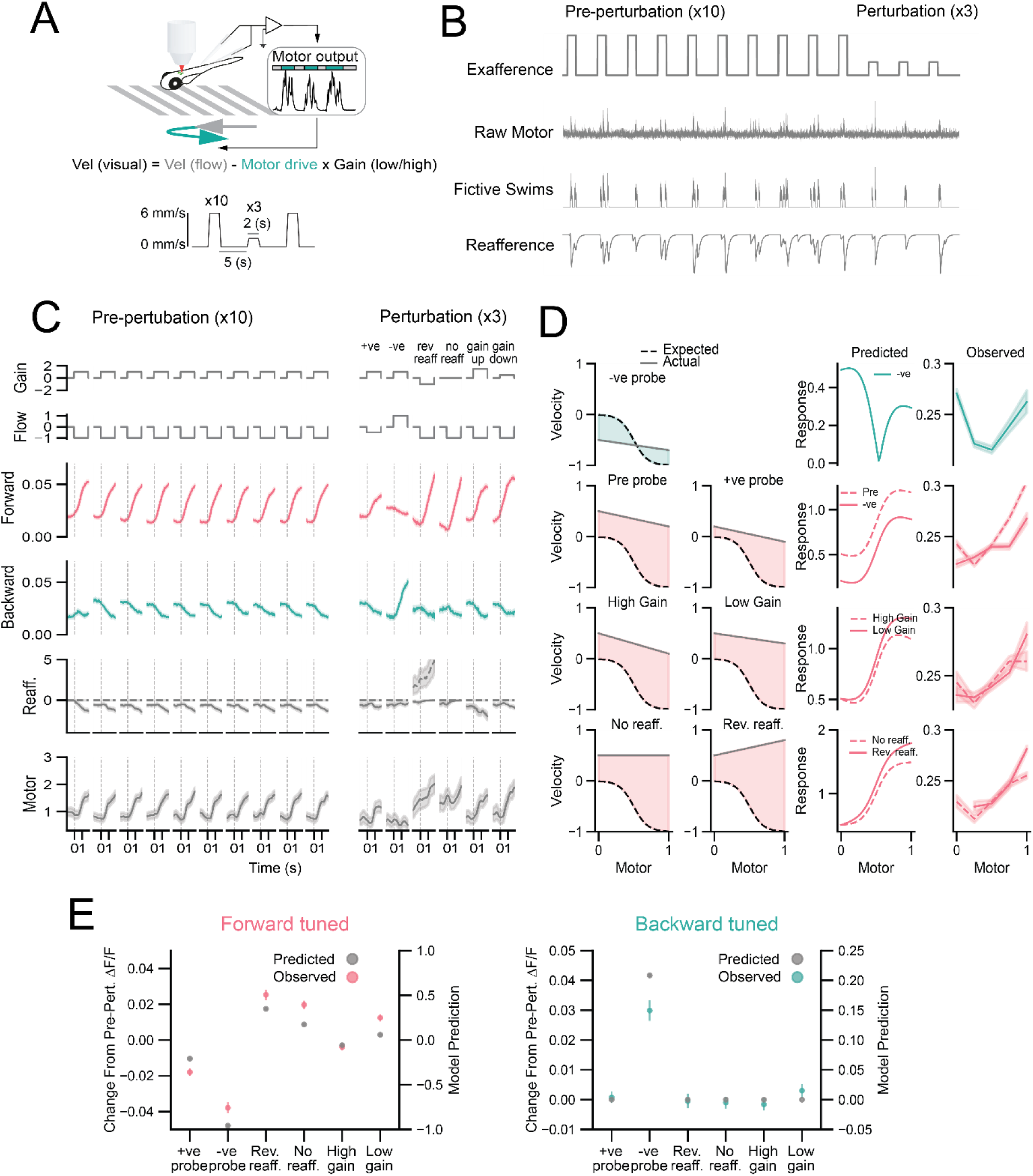
IO neurons signal the discrepancy between actual and expected feedback during closed-loop perturbations. A) Closed-loop virtual reality setup where electrically recorded fictive swims drive the visual scene according to a gain parameter. B) Example trial structure; ten pre-perturbation periods of forward moving gratings followed by 3 repeats of a perturbation, such as reduction in the velocity of the exafference. Raw motor was recorded from the zebrafish tail; fictive swims were extracted and the reafference signal was convolved with a GCaMP6f kernel. C) Forward-tuned and backward tuned responses pre-perturbation and during perturbations (+ve probe, -ve probe, reverse reafference, no reafference, gain up and gain down. D) Graphical hypotheses and predicted responses for each perturbation versus observed IO responses as a function of swim strength (Motor). E) Left: Comparison of perturbation-induced IO responses relative to pre-perturbation for forward-tuned neurons. Right: Comparison of backward-tuned IO responses relative to pre-perturbation.

Backward-tuned neurons only significantly responded to a change in the backward movement of the visual field (-ve probe), whereas forward-tuned neurons robustly reflected perturbations in reafference during forward-moving visual scenes (Fig 2E). Relative to the pre-condition, post-condition forward responses showed stimulus-specific modulation; responses increased for gain-down, no-reafference, and reverse-reafference probes (β = 0.012–0.025, all p < 0.001), decreased for negative and positive probes (β = −0.038 and −0.018, respectively, both p < 0.001), and did not differ significantly for gain-up probes (p = 0.105). For backward-tuned neurons, post-condition responses showed a selective increase only for negative probes (β = 0.030, p < 0.001), with no significant post–pre changes for any other stimulus (p ≥ 0.27). This demonstrated that IO responses scaled proportional to sensory feedback perturbation. IO neurons are extensively electrically coupled by gap junctions, which allow different degrees of synchronization between neurons that can be modulated by synaptic input within IO dendritic glomeruli e.g., ^11^. We therefore sought to address whether changes in IO population responses, and consequently the error signal, are defined by an increase in activity of IO neurons already active during pre-perturbation or instead defined by an increased recruitment of inactive neurons, for example via gap-junction mediated synchronization. By binning responses of forward-tuned neurons according to activity in pre-perturbation tests, we observed that the most active neurons were the first to increase activity during perturbations; however, more extreme perturbations produced increased responses of previously inactive neurons (Fig S1A-C). This implied that in low-error paradigms, error signals are conveyed by a small subpopulation of neurons, whereas high-error paradigms lead to increased recruitment of IO neurons. We observed little effect on future behavior after each perturbation (Fig S2) except for significantly increased latency to and amplitude of the first swim on the next trial after the negative probe (-ve) perturbation (Fig S2C, D).

Ablating the IO abolished this effect and demonstrated that it was IO dependent (Fig S2C, D).

### Inferior olive subpopulation responses can be recapitulated by error encoding model

We next formalized our hypothesis as a parameterized model, r ≈ ∣v−f(m)∣ where r is the IO neuron response, v is the actual visual velocity experienced by the fish and f(m) is a 4-degree polynomial representing expected visual velocity as a function of swim motor strength. We first fit this model to IO responses from open-loop moving dots experiments (see Fig 1), deriving an estimate of sensory expectation for different swim strengths (Fig 3B). From the derived polynomial curve, we could predict neuronal activity for a given swim power as proportional to the discrepancy between the actual velocity (VTrue) and the expected velocity (VExp). This produced predictions (Fig 3B, middle) of forward and backward-tuned IO responses that were highly or moderately correlated, respectively, with experimental data from trials used to fit the model and on trials held-out for validation (Fig 3B, right panel). In closed-loop experiments where the direction of visual flow was inverted (Fig 3C), we observed moderate correlation between model predictions and actual forward and backward-tuned responses (Fig 3D).

**Figure 3.**
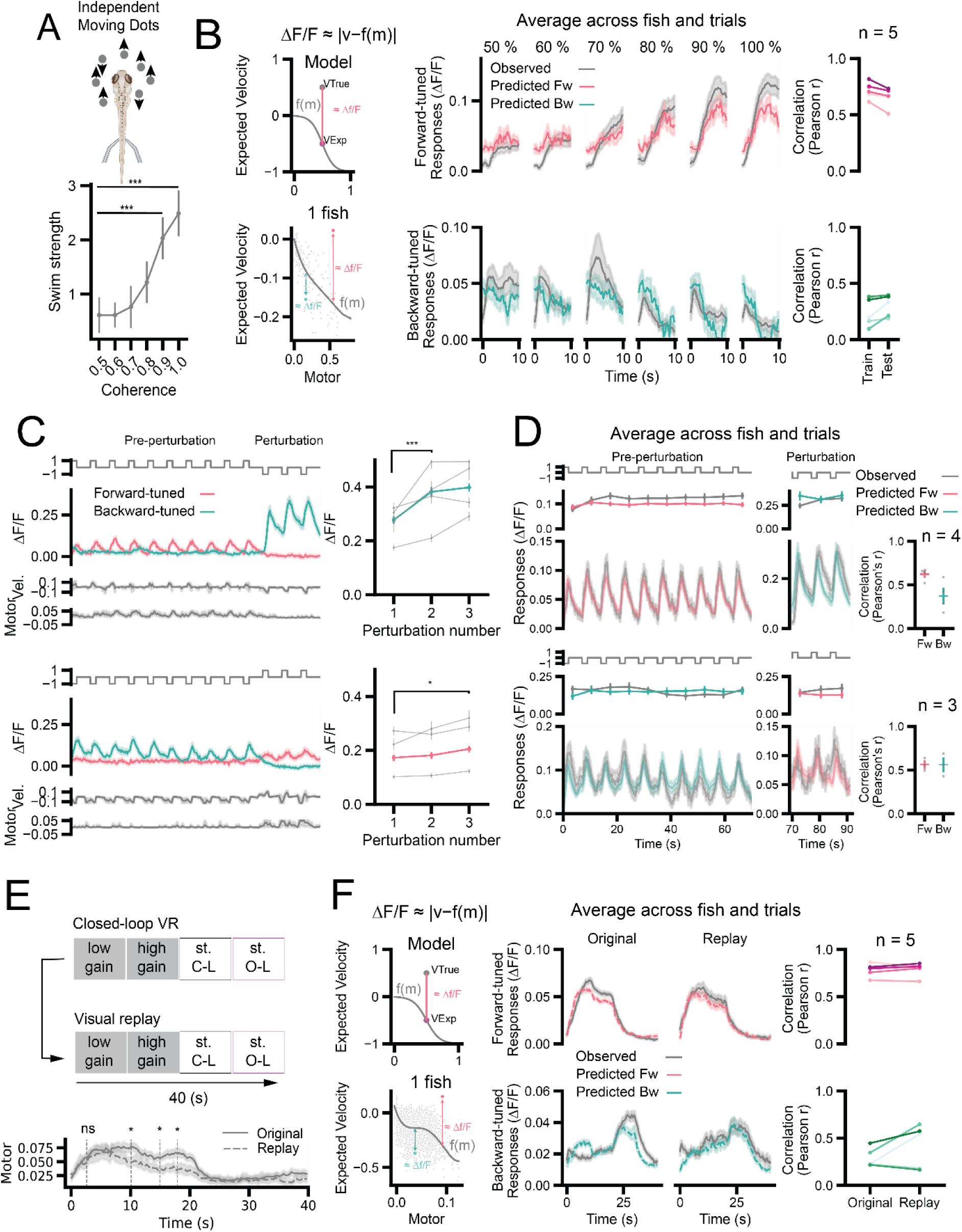
IO responses can be predicted from an approximated visual expectation. A) A series of dots moving in the same or opposite directions were presented to the fish. Swim strength as a function of movement coherence. B) Left panel: Graphical representation of error model and estimated visual velocity relationship (polynomial curve, top) with motor for one fish (bottom). Middle panel: Responses and model predictions for forward-tuned (top) and backward-tuned IO neurons at different dot movement coherence (50%-100%). Right panel: Pearson’s correlations between IO responses and model predictions for train trials used to fit the model and test trials ‘held-out’ from model fitting. C) Left panel: Repetitive visual motion in the forwards (top) or backwards (bottom) direction followed by a sudden change in visual motion direction. Right panel: Backward-tuned (top) and forward-tuned (bottom) IO responses as a function of perturbation repetition. D) Left panel: Direction-tuned IO responses and model predictions during pre-perturbation motion and perturbation motion. Right panel: Pearson’s correlations for forward (Fw) and backward (Bw) -tuned model predictions during both experiments. E) A closed-loop virtual reality gain change trial where the visual scene is replayed in replay trials. Trial order is low gain, high gain, stationary closed-loop and stationary open-loop. Bottom: A comparison of swim power in original and replay trials. F) Left panel: The formalized model where IO response (ΔF/F) is proportional to the absolute difference between actual velocity (VTrue) and the expected velocity (VExp) found on the f(m) curve (top). An example fish showing a polynomial f(m) can be approximated by fitting the model to forward-tuned neuron responses (bottom). This expectation f(m) can then be used to predict forward and backward tuned IO responses. Middle panel: Forward and backward-tuned IO responses (gray) and model predictions (colored) during the original trials and replay trials. Right panel: Pearson r correlation scores for comparison of model prediction and actual responses for each fish and original and replay trials.

However, we also observed significant facilitation in forwards- and backwards-tuned responses and habituation in backwards-tuned responses (Fig 3C) that was not fully captured by this model alone (facilitation: mixed-effects model, forwards-tuned, 1 vs 3 pulses, β =0.022, p <0.01, backwards-tuned, 1 vs 3 pulses β = 0.122, p<0.001; 2 vs 3 pulse, no difference, Tukey HSD, p = 0.59). We next performed a set of longer (40 s) closed-loop experiments where fish experienced feedback gain changes at 10 second intervals during forwards-triggered optomotor response (OMR) swimming, inducing the fish to adapt its swim power to stabilize itself in space ^4^. We then repeated these trials, replaying the visual stimulus from the original trial back to the fish (Fig 3E). Original trials exhibited significantly elevated motor activity at later time points compared to replay trials (mixed-effects model; motor × time interactions: β = 0.033–0.052, *p* ≤ 0.011), effectively decoupling motor behavior from sensory stimulus. This allowed us to fit our model to the original trials to extract the expected velocity curve, f(m), and test on the replay trials where behavior is the uncontrolled variable (Fig 3F top left). We learned an estimate of the expectation curve (f(m)) as before, by fitting the parameters to forward-tuned IO activity during the original trials (Fig 3F bottom left). This expectation curve, despite being derived during head-fixed virtual reality experiments, closely resembled the inverse sigmoid-like expectation curve found in freely swimming zebrafish larvae behavior during the same behavioral task (Fig 1A). From this, we were able to reproduce the response of forward-tuned IO neurons in the original trials and, importantly, also in the unseen test replay trials with moderate correlation (Fig 3F, top row). We then asked if this estimate of the expectation could be used to reproduce backward-tuned IO activity without the model having been fit to those responses. Promisingly, we were able to reproduce backward-tuned IO activity in the original and replay trials with moderate correlation for most fish suggesting that IO activity for opposing directions can be derived as the sensory discrepancy from a single estimate of internal expectation (Fig 3F, bottom row).

### Excitatory input to the IO is tuned to the direction of visual motion

We next asked what input drives error-encoding IO activity, first looking for pure motor input to the IO (Fig 4A). We recorded glutamatergic input into the IO reported by iGluSnFr ^18^ expressed in IO neurons (vGluT2+) and revealed few glutamate events in the IO before swim bouts but some infrequent events after swim bouts in the absence of visual motion (Fig 4B). Instead, we found that glutamatergic input was directionally tuned to the motion of the visual stimulus (Fig 4C, D), mirroring the spatial organization of the direction-tuning of the IO calcium responses (Fig 1B, right panel). Sensory-tuned glutamate events had a rightward-shifted cumulative distribution relative to motor events, indicating greater reliability and recruitment of sensory inputs than purely-motor inputs (Fig 4E). However, during closed-loop OMR (Fig 4F) and repeated perturbation (Fig 4G) experiments, we observed forwards- and backwards-tuned responses reminiscent to those in calcium imaging experiments (Fig 3C, F). We also observed facilitation of glutamate responses during directional motion changes suggesting that this phenomenon occurs upstream of the IO or at the pre-IO synapse (mixed-effects model; 2^nd^ pulse: β = 0.156; 3^rd^ pulse: β = 0.271; both *p* < 0.001 relative to 1^st^ pulse) (Fig 4H). The similarity of calcium and glutamate responses raised an interesting question; does the IO inherit error information upstream?

**Figure 4.**
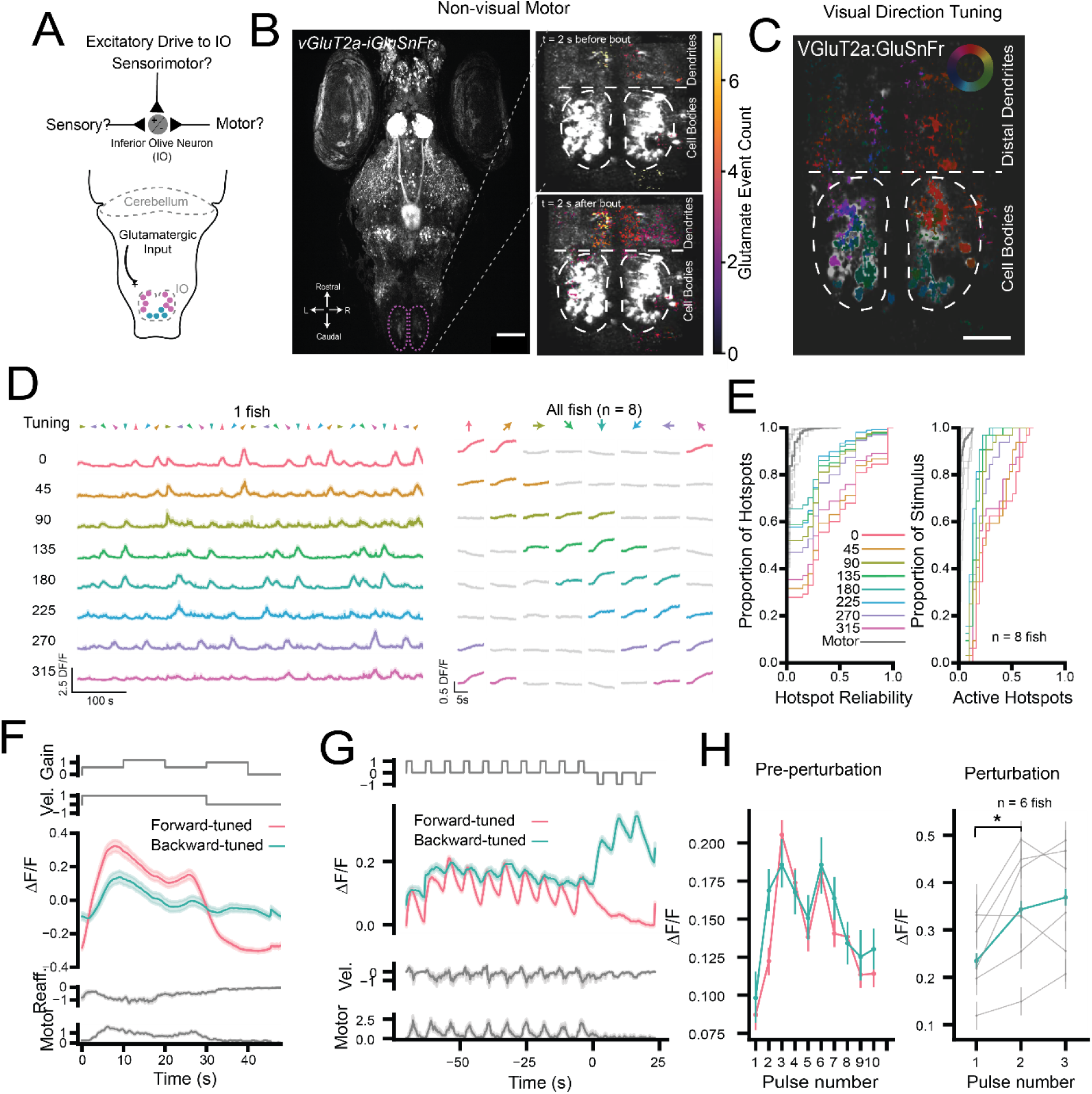
IO input is directionally tuned and mirrors calcium responses. A) Schematic showing glutamatergic input to the IO. B) Left panel: whole brain image from vGluT2a-iGluSnFr showing expression in inferior olive. Right panel: An example IO showing the total motor-related glutamate events in the dendrites and cell bodies of IO neurons before and after swim bouts. C) Spatial organization of directional tuning of glutamatergic input in the IO. D) Left panel: Average traces for glutamate events from one fish binned by directional preference. Right panel: Average direction stimulus responses for glutamate events binned by directional preference across all fish. E) Cumulative distributions showing the reliability and proportion of active direction-selective hotspots compared to motor hotspots. F) Average forward- and backward tuned glutamate responses across fish during closed-loop OMR with feedback changes (average reafference and motor in gray). G) Average forward- and backward tuned glutamate responses during ten consecutive 2 second pulses of forward visual motion followed by three pulses of backward visual motion. H) Left: Maximum responses for forward- and backward-tuned glutamate signals for each pre-perturbation pulse, averaged across fish. Right: Maximum responses for backwards-tuned input for each perturbation pulse, averaged across fish and individual fish shown in gray. Scale bars: 100µm in B, 50µm in C.

### IO neurons receive prediction error signals in excitatory input and selectively gate transmission to the cerebellum

To test whether IO input reflected sensory errors (Fig 5A, C), we simultaneously recorded glutamatergic input (vGluT2a-iGluSnFr) to the IO as well as glutamate release or spillover from climbing fiber terminals in the cerebellum (Fig 5B) during visual motion at a constant velocity in open loop in either the forwards or backwards direction (Fig 5D). Parallel fibers are vGluT2a-negative ^23,24^ allowing us to exclusively monitor glutamate release from vGluT2a-positive climbing fibers. This identified that both IO input and output were modulated by swim strength (Fig 5E, F), rejecting the null hypothesis that the IO input should be constant if the IO was directly contrasting actual constant and expected visual velocity to produce a prediction error (Fig 5C). Instead, this showed that IO neurons directly receive pre-computed error-like signatures as motor-modulated signals, originating from other brain regions. To evaluate this in closed-loop in response to swimming behavior, we performed random reafference perturbations on a bout-by-bout basis, sampling IO responses across a range of bout-triggered visual scene changes and swimming motor commands (Fig 5G). We mapped iGluSnFr responses for forward- and backward-tuned IO inputs and outputs in visuo-motor space and observed non-uniform receptive fields for both neuron types (Fig 5H). Forward-tuned input increased non-linearly as a function of forward visual velocity and motor swim signals (Fig 5I). This was broadly mirrored by the receptive field for forward-tuned output; however, peak activity was sparser and suggested a filtering effect between IO input and output (Fig 5I). Backward-tuned input displayed multiple peaks surrounding a valley whilst backward-tuned output represented a higher contrast version of the input (Fig 5J). IO responses were generally larger for regions of the visuomotor space where swim bouts were less likely to occur. We assumed that direction tuning is maintained from IO input to output and analyzed the coherence between averaged directional (forwards/backwards) input and output signals. This represents the frequency dependent transfer of information from input to output and revealed a prominent peak coherence at lower frequencies and attenuation at relatively higher frequencies (within the imaging sampling frequency of 4.5Hz, mixed-effects model; 0.3 Hz vs frequencies ≥0.9 Hz (all p ≤ 0.047)) (Fig 5K). This was consistent with the inferior olive acting as a low-pass filter.

**Figure 5.**
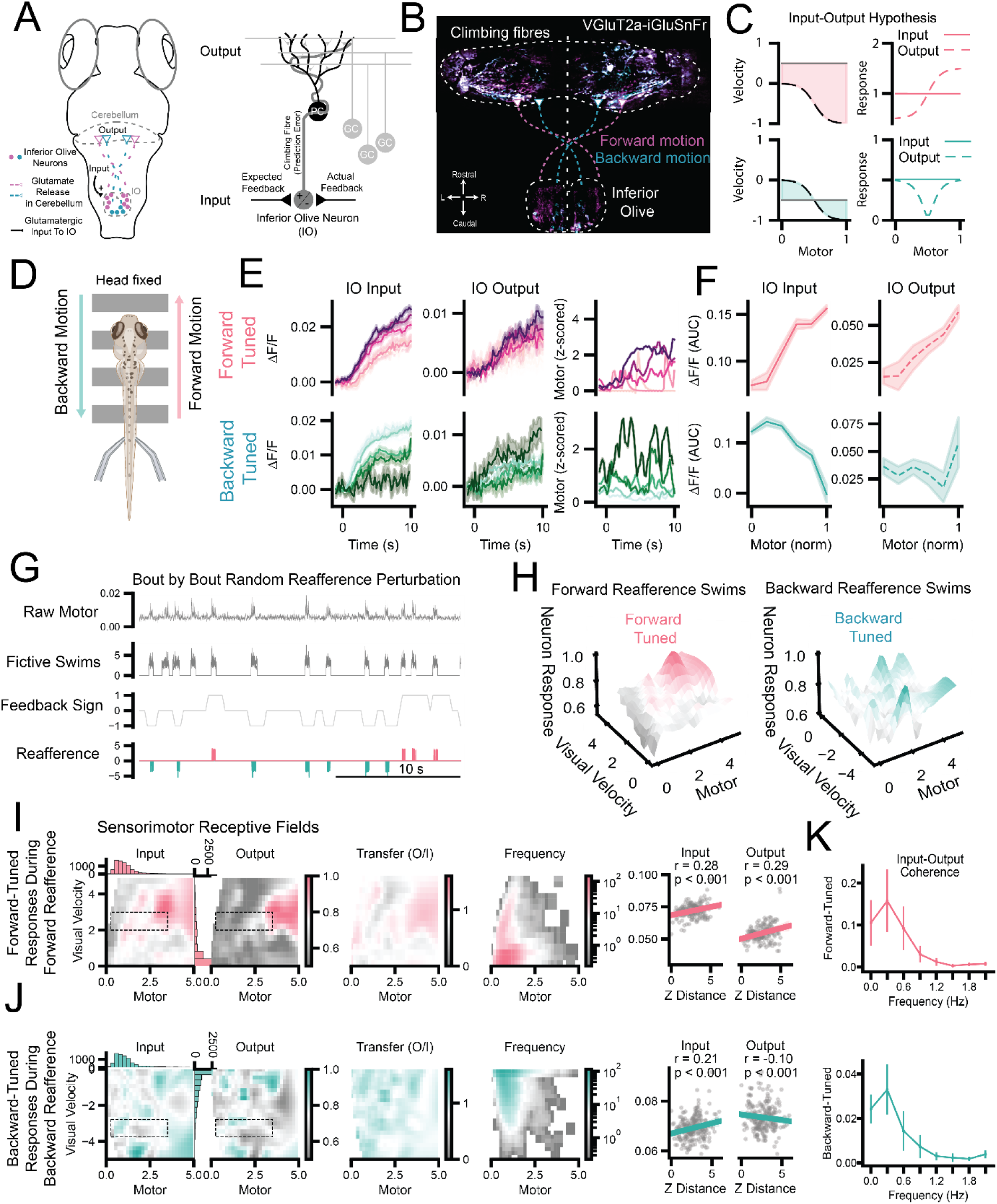
Excitatory input encodes prediction errors and IO neurons selectively transmit these to the cerebellum IO. A) Left: Schematic describing anatomical input to the IO and climbing fiber output to the cerebellum in zebrafish. Right: Olivo-cerebellar circuit detailing the comparator hypothesis for the IO in computing prediction errors. B) GluSnFr fluorescence in cell bodies in the IO and climbing fibers in the cerebellum – color-coded by direction of visual motion. C) The error hypothesis for neuronal responses and input to the IO. D) Zebrafish larvae were head fixed and moving gratings were projected underneath the fish. E) Forward and backward tuned glutamate hotspots binned by swim strength at the input to IO and at the climbing fiber output from the IO to the cerebellum. F) Relationship between swim strength (Motor) and iGluSnFr response at IO input and IO output. G) An example trial from one fish showing random reafference perturbation after each fictive swim. H) 3D surface plots showing forward and backward tuned glutamatergic input to the IO during forward reafference and backward reafference swims, respectively. I,J) Left: 2D plot showing receptive field of glutamatergic input and output. Right: Linear regression of glutamate response as a function of distance in z-standardized visuomotor space. K) Coherence plots representing transfer of information from input to output as a function of frequency (note 0.0 Hz represents steady state and the correlation of the average of the signals).

### IO neurons shape learning during longer-term behavioral adaptation

Our results suggest that IO neurons filtered, rather than computed, prediction errors, acting as low-pass filter; we therefore examined the necessity of the IO for instructive cerebellar-dependent learning at the relevant timescales. Our findings here, as well as previous work ^4^, have indicated that the IO is essential for some short-term behavioral adaptations in zebrafish; however, the role of the IO in longer-term behavioral adaptations is relatively unexplored. Here, we define short-term for zebrafish on the order of seconds and longer-term on the order of minutes or hours. We analyzed IO dynamics during two established behavioral tasks that exhibit longer-term changes in behavior as the zebrafish tries to achieve the ethological goal of stabilizing itself in space ^6^. The first task subjected the fish to 40-second-long forward moving gratings where the closed-loop feedback gain varies every 10 seconds, challenging the fish to adapt its swim strength and swim bout frequency to minimize displacement (Fig 6A). We measured the final displacement of the fish at the end of each trial, fitting a linear model to each fish’s final displacement over 30 trials. We classified fish as initial overshooters or undershooters if the y-intercept was greater or less than 0.5 SD, respectively, and found that overshooters displayed a negative learning slope thereby reducing overshooting across trials (Fig 6A). The opposite for undershooters was also observed, indicating that the fish’s initial performance dictated the learning direction and that both groups improved, minimizing displacement over trials (Fig 6A). There was a strong correlation between the displacement of the previous and current trial, indicating that zebrafish used an incremental learning strategy, making small steps towards improving performance (Fig S3A). Direction tuning of IO neurons was computed as before (Fig S4A-C) and forward- and backward-tuned IO responses varied during the trial as the gain changed (Fig 6B) and decreased over trials (Fig 6C, S3B). This suggested that as fish improved at the task, performance error decreased and thus error-related signals in the IO also decreased (Fig 6C). Interestingly, the activation slope of neuron responses was graded to the degree that neuron direction preference was off the axis of the visual stimulus direction. This suggested that IO neurons integrate errors at time constants according to the projection of the error on the neuron’s direction preference (Fig S4 D). We selectively ablated forwards and backwards-tuned IO populations and assessed the short-term and long-term performance of fish during the task. Despite a general increase in swim power, neither targeted ablation impaired the ability of fish to adapt their motor output to the changing feedback gain, with short-term adaptation preserved (Fig 6D). However, ablation of the entire IO, consistent with prior work, did significantly impair short-term adaptation (Fig S3C). Swim duration on the first bout was slightly but significantly increased after forward-tuned IO ablation only (Wilcoxon, p < 0.05), and no changes in latency to the first bout were observed (Fig 6F). When analyzing longer-term performance, laser ablating forward-tuned neurons significantly altered the ability of overshooters to converge (mixed-effects, β= 0.06, p<0.001), whereas ablating backward-tuned neurons conversely, but not to significance, perturbed the convergence of undershooters (Fig 6G). This suggested that these oppositely direction-tuned populations of IO neurons help implement the direction of incremental learning over longer-term timescales.

**Figure 6.**
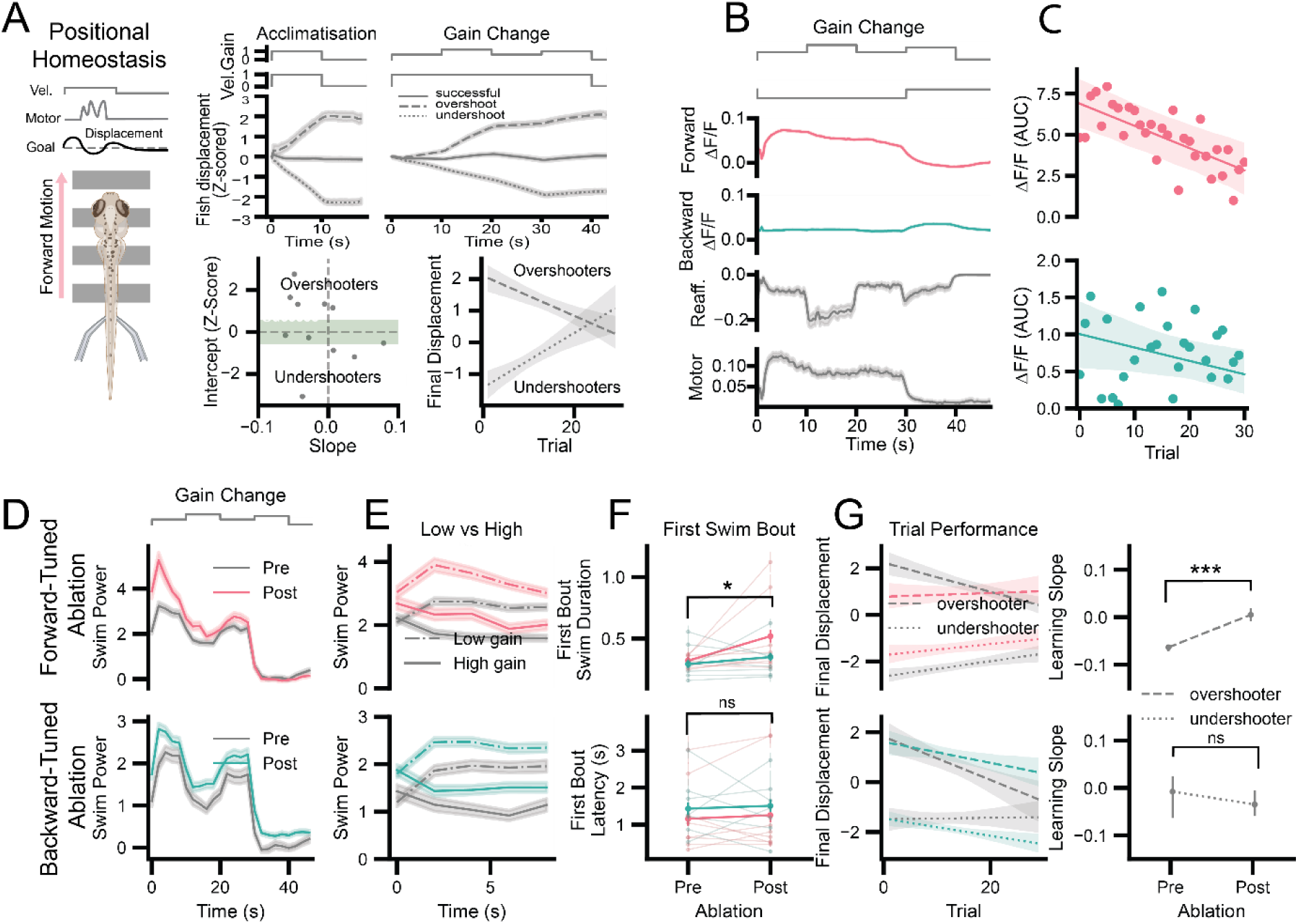
IO neurons drive longer-term behavioral adaptation after perturbation. A) A positional homeostasis task where fish swim in response to forward-moving visual gratings projected beneath the fish. Trials were characterized as overshoots, undershoots or successful according to the final displacement at the end of each trial. Fish were classified as undershooters or overshooters by a linear regression of final displacement by trial where a positive y-intercept defines a fish that predominantly overshoots in the first trials, and vice versa for undershooters. B) Average IO activity for forward-tuned and backward-tuned neurons during an average trial. C) Linear regression of average forward- and backward-tuned IO neuron activity over trials as fish improve. D) Swim power during the average positional homeostasis trial before and after ablation of forward-tuned (top) and backward-tuned (bottom) neurons. E) Swim power during the low-gain and high-gain part of the average positional-homeostasis trial before and after ablation of forward-tuned (top) and backward-tuned (bottom) neurons. F) Left: Duration of (top) and latency (bottom) to the first swim bout before and after ablation of forward-tuned (pink) and backward-tuned (green). G) Linear regression fit of trial performance (final displacement) for overshooters (dashed-line) and undershooters (dotted-line) before (gray) and after ablation (colored). Right: Quantification of these results across trials and fish (n=14).

## Discussion

This study addressed a longstanding question in cerebellar theory: whether the inferior olive (IO) computes prediction errors locally or relays signals that are already structured by upstream processing. By combining population calcium imaging with measurements of glutamatergic input and climbing fiber output during controlled visuomotor perturbations, we were able to compare synaptic drive to IO neurons with the signals transmitted to the cerebellum under identical behavioral conditions.

Three main findings emerged. First, IO activity scaled with discrepancies between expected and experienced sensory feedback across open- and closed-loop contexts. Second, excitatory input to the IO already contained structured, directionally-tuned and motor-dependent components consistent with error-like information. Third, transmission from IO to cerebellum was selective: output represented a filtered version of input, with preferential transfer of lower-frequency components and contrast enhancement in specific regions of visuo-motor space. Together, these results indicate that prediction error-like structure does not arise solely within the IO but is substantially shaped upstream and then transformed by the olive before reaching the cerebellum.

### Locus of error computation

Classical models often assign the IO a comparator role in which expected and actual feedback are contrasted to generate a teaching signal such as in the vestibulo-ocular reflex (VOR) ^25^. Our measurements of glutamatergic input challenge a strict version of this view. For instance, under conditions where visual velocity was constant, excitatory drive to the IO varied with swim strength. Thus, motor-dependent structure is already present in afferent input before olivary processing.

This observation supports models in which upstream circuits participate in constructing discrepancy signals that are subsequently routed through the IO, as has been described for the OKR ^25^. In this context at least, the olive appears not as the origin of error computation and more as a node that selects, synchronizes, and formats instructive information for cerebellar plasticity.

### Transformation between input and output

Although input already exhibited error-like structure, output was not a simple copy. Simultaneous recordings revealed sparser activation, enhanced contrast, and strong attenuation of higher-frequency components. Coherence analyses indicated preferential transmission at slow timescales, consistent with IO neurons transforming high-frequency sensory-errors input into a low-frequency instructive climbing fiber teaching signal ^26^.These properties suggest that the IO implements a gating or filtering operation ^11^. Rather than computing errors from scratch, the IO may regulate when and how upstream discrepancy signals are allowed to influence cerebellar learning. This regulation could arise from intrinsic oscillatory mechanisms linking error signaling to the temporal structure of behavior, intra-IO electric coupling, and inhibitory control from nuclei including cerebellum ^27^. Interestingly, forward- and backward-tuned pathways differed in how strongly output mirrored input, implying asymmetric input or additional sources of excitation or inhibition for particular channels. Determining the anatomical origin of these subpopulation-specific differences will require future circuit mapping.

### The role of glutamate in the IO

Glutamate is the major excitatory neurotransmitter in the IO, with primary projections (in mammals) from the motor cortex, red nucleus, mesodiencephalic junction and deep cerebellar nuclei ^28^. Zebrafish lack a motor cortex, and glutamatergic projections to the IO are unmapped; however, homologs for the red nucleus ^29^, deep cerebellar nuclei (eurydendroid cells in zebrafish ^30^), and mesodiencephalic junction structure exist and these likely represent the source of excitatory sensorimotor input to the IO. But what is the effect of glutamate on IO function? Glutamatergic inputs either synapse within glomeruli to influence coupling of dendro-dendritic electrical synapses and synchrony of input ^31^, or terminate more directly onto dendrites and somata to influence IO neuronal firing. In the case of the latter, glutamate weakly increases firing frequency by around 2-3Hz ^32^ and spontaneous activity in the IO does not require active excitation. Thus, it’s important to question what underlies the error signal recorded by calcium reporters – subtle increases in firing frequency, changes in the amplitude and number of spikelets of the waveform of the action potential or changes in the proportion of neurons synchronized in activity. By analyzing the contribution of recruitment of neurons and changes in gain of neurons already recruited we propose that the underlying calcium signal predominantly reflects an increase in per-neuron gain (frequency or waveform) in low-error paradigms and an increase in recruitment of silent IO neurons in high-error paradigms. Additionally, stimulation of excitatory projections to the IO results in a phase-resetting of sub-threshold oscillations ^11^, adjusting the future spiking pattern of the neuron, where action potentials and their waveforms are likely to ride and be modulated by the peaks of the phase-adjusted oscillations.

### Role in short- and long-term adaptation

The dichotomy and debate of the role of the inferior olive in the central nervous system centers on whether the IO directly modulates motor behavior or induces plasticity and learning in the cerebellum^33^. Here, we find that these are not mutually exclusive and present evidence for influence on motor control in the short term as well as in cerebellar-dependent longer-term learning. Ablation experiments demonstrate that IO activity contributes to behavioral adaptation across multiple timescales. Global IO removal impaired both rapid adjustments to perturbations and slower convergence across trials, confirming an instructive role in motor adaptation. In contrast, selective ablation of directionally tuned subpopulations primarily altered longer-term learning while sparing short-term compensation. This distinction suggests that distributed IO ensembles guide gradual plasticity in cerebellar circuits^34–38^, whereas immediate motor adjustments depend on broader population recruitment or parallel pathways^39–43^.

### Temporal dynamics beyond the discrepancy model

Repeated perturbations induced facilitation and habituation that were not fully captured by the discrepancy model. Such dynamics may reflect saliency-dependent transmission, changes in inhibitory balance ^6^, or presynaptic mechanisms regulating glutamate release. Because facilitation was already present in excitatory input, at least part of this computation likely occurs upstream of the IO. Future studies will reveal the mechanisms underpinning these dynamics, which could include a combination of neuromodulation, presynaptic facilitation, or intrinsic neuronal properties.

### Directional organization and population coding

Excitatory input mirrored the topography of IO directional tuning, suggesting that preference is inherited rather than synthesized locally. Although individual neurons primarily signaled the magnitude of mismatch, the relative activity between the oppositely tuned neuronal populations preserves directional information, allowing cerebellar circuits to infer how behavior should be modified. Because neurons with different preferred directions are driven in proportion to how strongly the discrepancy aligns with their tuning, the population collectively provides information that can guide distributed, gradient-like adjustments in behavior implemented by cerebellar circuits.

### Future directions

Our results implicate the IO in both the timing of initiation of behavior as well as continuous adjustment of motor output. This supports prior work demonstrating that IO activity is predictive of future swimming onset ^6^ . However, in zebrafish, acute motor adaptation does not appear to be cerebellum dependent ^5^; therefore, IO-mediated adjustment of motor behavior in the short-term in zebrafish must occur via another pathway. Since in mammals IO neurons are known to project collaterals directly to DCN neurons ^28^, it will be interesting to determine if an equivalent pathway exists and has functional implications for reflex-like control of movement in zebrafish.

### Limitations

Here, we leveraged the optical accessibility of zebrafish to test fundamental theories of IO function at the rarely observable population level^4,6,17^. However, several experimental constraints limit our ability to draw conclusions. Calcium signals provide indirect measures of spiking and cannot resolve spikelets or subthreshold dynamics that are central to olivary physiology ^16^. iGluSnFR reports extracellular glutamate rather than synaptic currents ^44^, and therefore may blur distinctions between direct excitation and spillover. In addition, our paradigms focused primarily on visual feedback, whereas vestibular and proprioceptive signals are also likely to contribute to natural error representations ^45,46^ .

### Conclusions

By directly comparing excitatory input to IO neurons with climbing fiber output during behavior, we provide evidence that error-like information is substantially organized before reaching the olive. The IO then reshapes this input through selective, frequency-dependent transmission, producing teaching signals suited for cerebellar plasticity. These findings refine the classical comparator framework and place the IO as a critical transformation stage within a distributed network for constructing instructive signals for motor learning and adaptation.

## STAR Methods

### Key resources table

| REAGENT or RESOURCE | SOURCE | IDENTIFIER |
| --- | --- | --- |
| Chemicals, peptides, and recombinant proteins |  |  |
| alpha-bungarotoxin | Alomone Labs | B-100 |
| Experimental models: Organisms/strains |  |  |
| Zebrafish:<br>Tg(elavl3:H2B-GCaMP6f) | Vladimirov et al., 2014 | N/A |
| Zebrafish:<br>Tg(vGluT2a:GAL4;UAS:iGluSnFR) | This study | N/A |
| Software and algorithms |  |  |
| ScanImage | Pologruto et al., 2003 |  |
| CalmAn | Giovannucci et al., 2019 |  |
| Bonsai | Lopes et al., 2015 |  |
| Custom Python scripts | This study |  |
| Deposited data |  |  |
| Processed data after CalmAn pipeline | This study |  |

### Animal Husbandry

All animal handling and experimental procedures conformed to the UK Animals (Scientific Procedures) Act (ASPA), 1986, and were approved by the UK Home Office and the Animal Welfare Ethics Committee (AWERB) of the University of St Andrews (Project licence numbers PP8440114 and PP4780998). All experiments were performed on larval zebrafish (Danio rerio) at 5-7 days post fertilization (dpf) reared in E3 medium at 28°C with a 14/10 h light-dark cycle.

### Transgenic lines

Experiments were conducted in two transgenic zebrafish lines: Tg(elavl3:H2B-GCaMP6f), expressing a nuclear-localized calcium indicator under the pan-neuronal elavl3 promoter, and crossed on casper or nacre backgrounds. The other, Tg(vGlut2a:GAL4;UAS:iGluSnFR), expressing the glutamate sensor iGluSnFR in glutamatergic neurons obtained from a cross between Tg(UAS-iGluSnFR, gift from Lagnado) and Tg(VGlut2a:GAL4) adults.

### Method details

#### Preparation of zebrafish for imaging

All imaging experiments were performed using 5-7 day old larvae. On the day of imaging larvae were immobilized by bath application of alpha-bungarotoxin dissolved in E3 medium (0.08 mg/ml, Alomone Labs, B-100), for approximately 10-20 minutes, then embedded in 1.7% low melting point agarose (Sigma Aldrich) in 35 mm petri dish, with the agarose around the tail removed to allow for attachment of suction electrodes for locomotor electrophysiology. Tail recordings were taken with a Differential AC Amplifier (AM-Systems). Zebrafish were mounted under a two-photon microscope with a 16x objective lens (Nikon, CF175 LWD). Visual scenes were projected at 60 fps using a projector, onto diffuse acrylic beneath the recording chamber using a mirror (Edmund Optics) and a red-pass filter ensuring shorter wavelengths did not corrupt the two-photon images.

#### Two-photon microscopy

A custom built two-photon microscope with Spark Lasers ALCOR femtosecond pulsed laser with fixed wavelength (920 nm) was used to monitor brain fluorescence. Green fluorescence was collected using a Nikon 16X CFI LWD Plan Fluorite Objective and detected using a PMT with a green bandpass filter. A resonant galvo-galvo scanner (Vidrio RMR), allowed x, y scanning and a piezo for focusing in z, as well as 3D online motion correction. During GCaMP imaging, the IO was scanned at either 3.07 Hz or 15 Hz, each volume consisting of 5 planes, 5 µm apart whilst iGluSnFr imaging was performed at 4.56 Hz. The electrophysiology and stimulus presentation were controlled through Bonsai, and the two-photon imaging was controlled through ScanImage. A NI-DAQ (National Instruments) was used for data acquisition and the Bonsai package Bonsai.DAQmx used for interfacing with the data acquisition and control hardware; volume clock triggers were acquired through this to synchronize imaging with stimulus presentation and electrophysiology. At the end of all experiments an overview of the whole fish brain was acquired by scanning the entire brain and imaging 250 planes each 1 µm apart reaching a total depth of 250µm to be used for anatomical brain registration of the IO.

#### Two-Photon laser ablations

Larvae expressing Huc:H2B GCamp6f were prepared for IO ablations using the same protocol used for calcium imaging experiments. For ablations involving forward-tuned or backward-tuned cell populations, visual gratings were given in these directions to identify the relevant populations. Individual cells were selected manually to determine the laser path for ablations. Using ScanImage Photostimulation (MATLAB) the laser beam power was set to 90% and a log spiral pattern was set for a duration of 50 milliseconds for each manually selected ROI with a repetition pattern of three. Lesions were confirmed visually by an increase in the fluorescence intensity of each cell.

#### Discrepancy model/ expected swim feedback

The formalized model:

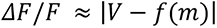

where the response of IO cells (ΔF/F) is proportional to the absolute difference between actual velocity (*V*), and the expected velocity *f*(*m*) defined as velocity expressed as a function of motor output. To determine the shape of *f*(*m*), velocity was plotted as a function of mean tail frequency (Hz) during forward OMR-induced swims in freely swimming larvae by fitting linear, log-linear and polynomial models to the data. Mean tail frequency was taken as a proxy for motor output. A 4-degree polynomial model produced the highest explanatory power for behavioral data (R^2^ = 0.47; Fig 1A) and was therefore used when fitting the model to forward-tuned IO GCaMP traces. The polynomial (f(m)) after fitting to forward-tuned IO population activity could then also be used to derive backward-tuned IO activity with the addition of a scaling parameter.

#### Open loop virtual reality experiments

##### Visual direction

To quantify direction tuning of individual IO neurons, we measured calcium or glutamate activity in response to moving square-wave gratings presented in open loop. The visual stimulus consisted of alternating red and black vertical bars (spatial period: 1 cm) that translated across the visual field at 0.6 cm s⁻¹ in eight directions (0°–315° in 45° increments), where 0° corresponded to forward motion (caudo-rostral). Direction tuning for each neuron was computed by taking the average of the vector sum across repeated presentation of directional stimulus.

#### Coherence/ moving dots

Visual scenes composed of randomly distributed dots moving at a constant velocity of 0.6 cm/s were projected beneath immobilized GCaMP6f-expressing zebrafish. To introduce varying levels of sensory uncertainty, the proportion of forward- and backward-moving dots was systematically altered across trials. Each trial contained one of six motion ratios: 50/50, 60/40, 70/30, 80/20, 90/10, or 100% forward-moving dots (Fig 1D-E, Fig 3A).

#### Closed loop virtual reality experiments

##### Perturbation experiments

To examine how IO neurons respond to sudden sensory perturbations, we employed perturbation experiments based on previously established stimulus protocols ^47^ in GCaMP6f- or iGluSnFR-expressing fish. Forward-moving visual stimuli were presented as 2 s closed-loop epochs interleaved with 5 s stationary periods. During closed-loop epochs, fictive swim bouts triggered visual feedback proportional to motor output (see OMR methods). Each trial comprised ten 2 s forward-motion pulses at 0.6 cm s⁻¹, followed by a perturbation sequence repeated up to three times (Fig 2, 4G). Inverse-perturbation trials consisted of backward-moving probe pulses (−0.6 cm s⁻¹) followed by the same perturbation sequence (Fig 3C–D). Perturbation stimuli consisted of 2 s probe epochs in which either the external visual scene or visual reafference was altered. Probe conditions included: (i) negative probe, in which the visual scene moved backward; (ii) positive probe, in which forward velocity was altered; (iii) gain-up and gain-down probes, which increased or decreased the strength of visual feedback; (iv) reverse reafference, in which the sign of visual feedback was inverted; (v) no-reafference probes, which mimicked open-loop conditions; and (vi) no-probe controls, in which there was no external motion but visual feedback was provided in response to swim bouts. Each trial was repeated up to ten times. Following baseline recordings, the inferior olive was bilaterally ablated in GCaMP6f cohorts using two-photon laser ablation (see Methods), and the full stimulus protocol was repeated after a 30 min recovery period (Supplementary Fig 2).

##### Reverse reafference perturbation experiments

To directly test the effect of random changes in visual reafference during ongoing closed-loop behavior, we extended the perturbation paradigm by presenting continuous forward-moving visual gratings in closed loop while randomly inverting the sign or gain of visual feedback after each swim bout (Fig 5G–K). This manipulation generated unexpected sensory consequences without changing the external stimulus.

##### OMR experiments

To measure short term motor adaptation, GCaMP6f- or iGluSnFR-expressing fish were presented with square gratings moving forward at a constant velocity of 0.6 cm s⁻¹ to evoke the optomotor response. Fish swam to counteract the apparent backward displacement of the visual scene. During fictive swimming, visual feedback was provided in closed loop such that the visual scene was momentarily accelerated backward in proportion to the detected swim bout, mimicking self-motion. Swim signals were processed online in Bonsai, and locomotor drive was quantified as the area under the rectified swim signal. To establish appropriate gain values, each fish underwent ∼20 acclimation trials in closed loop, during which a medium gain was manually adjusted to yield stable swimming. Low and high gains were then defined as at least two units below and above this baseline, respectively. Fish were subsequently exposed to ∼30 trials in which the visual scene moved forward continuously while gain alternated every 10 s according to the sequence: low–high–low–high–zero. To assess the contribution of direction-tuned IO populations to gain adaptation, subsets of fish underwent targeted two-photon ablation of either forward-tuned or backward-tuned IO neurons, after which the full gain-adaptation protocol was repeated (Fig 6).

##### OMR-Replay experiments

To dissociate sensory responses from motor commands, we performed replay experiments in GCaMP6f larvae (Ahrens et al., ref). Fish first experienced a gain-adaptation trial in closed loop as described above, consisting of 10 s of forward motion at low gain, 10 s at high gain, followed by 10 s of stationary gratings in closed loop (allowing swimming with feedback), and finally 10 s of stationary gratings in open loop. The entire 40 s visual scene was recorded and immediately replayed to the same fish in open loop (Fig 3E), such that visual input was identical but uncoupled from motor output. Each closed-loop/replay pair was repeated up to 12 times. Fish that failed to adapt to gain changes were excluded from analysis.

##### Image analysis

Inferior olive cells and their activity traces were extracted from high-quality components from the imaging data based on temporal and spatial criteria through CaImAn. Fluorescence signals from these cells were extracted as delta F/F values. Raw electrophysiology signals from the left and right tail were first convolved to a GCaMP6f or iGluSnFR kernel and resampled to match the imaging rate. All data analysis was conducted in Python.

##### Data analysis

Recordings were motion corrected and preprocessed using the CaImAn python package^48^. Inferior olive cells and their activity traces were extracted using non-negative matrix factorization and selected using temporal and spatial quality control criteria. Fluorescence signals from these cells were extracted as delta F/F values. Raw electrophysiology signals from the left and right tail were first convolved using a GCaMP6f or iGluSnFR kernel and resampled to match the imaging rate. All data analysis was conducted in Python and is available alongside. Statistical methods are reported in text, primarily using linear mixed effects models to account for individual fish variation.

## Data Accessibility

All data are available from the accompanying Zenodo repository at 10.5281/zenodo.18961438.

## Acknowledgments

We would like to acknowledge Jacqueline MacPherson, Michael Kinnear, Angus Aitken, Amy Dorward, and Thomas Powell from the Psychology Workshop for their technical support. We would also like to thank Holly Armstrong and Joe Chapman and technical staff from the Scottish Ocean Institute for zebrafish husbandry support and Leon Lagnado and Kate Hampden-Smith for the UAS:GluSnFr transgenic line.

## Supplementary Figures

**Figure S1.**
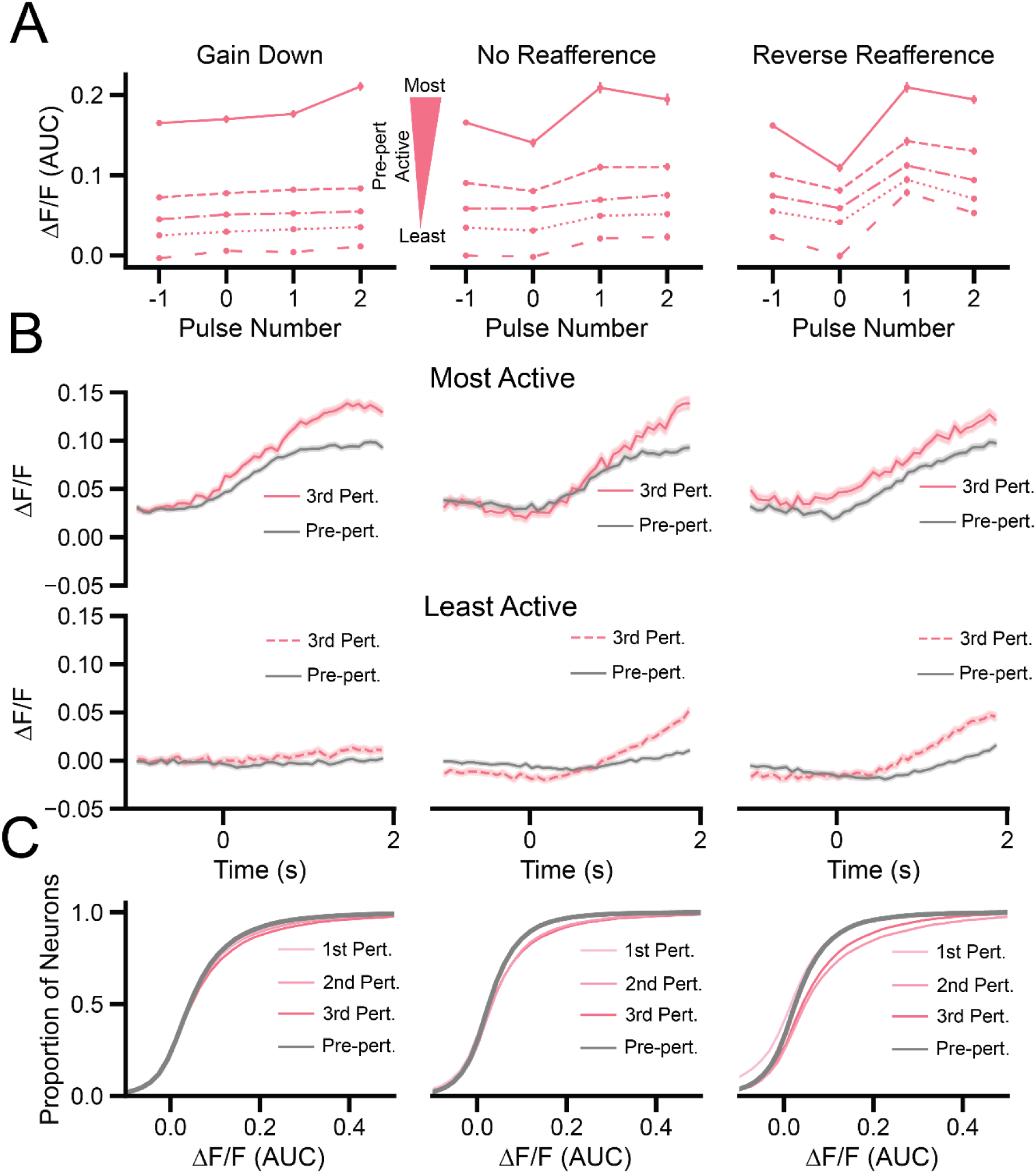
Increased gain of pre-active neurons and increased recruitment of inactive neurons contribute to IO response. A) Forward-tuned responses for perturbation compared to pre-perturbation binned by activity in the pre-perturbation control, ranging from most active (last quintile, solid line) to least active (first quintile, spaced dashed line). B) Upper row: traces of most active (last quintile) forward-tuned neurons compared to pre-perturbation average. Bottom row: traces of least active (first quintile) forward-tuned neurons compared to pre-perturbation (gray). C) Cumulative distribution plots comparing shifts in distribution for forward-tuned neuron responses during repeated perturbations, compared to pre-perturbation (gray).

**Figure S2.**
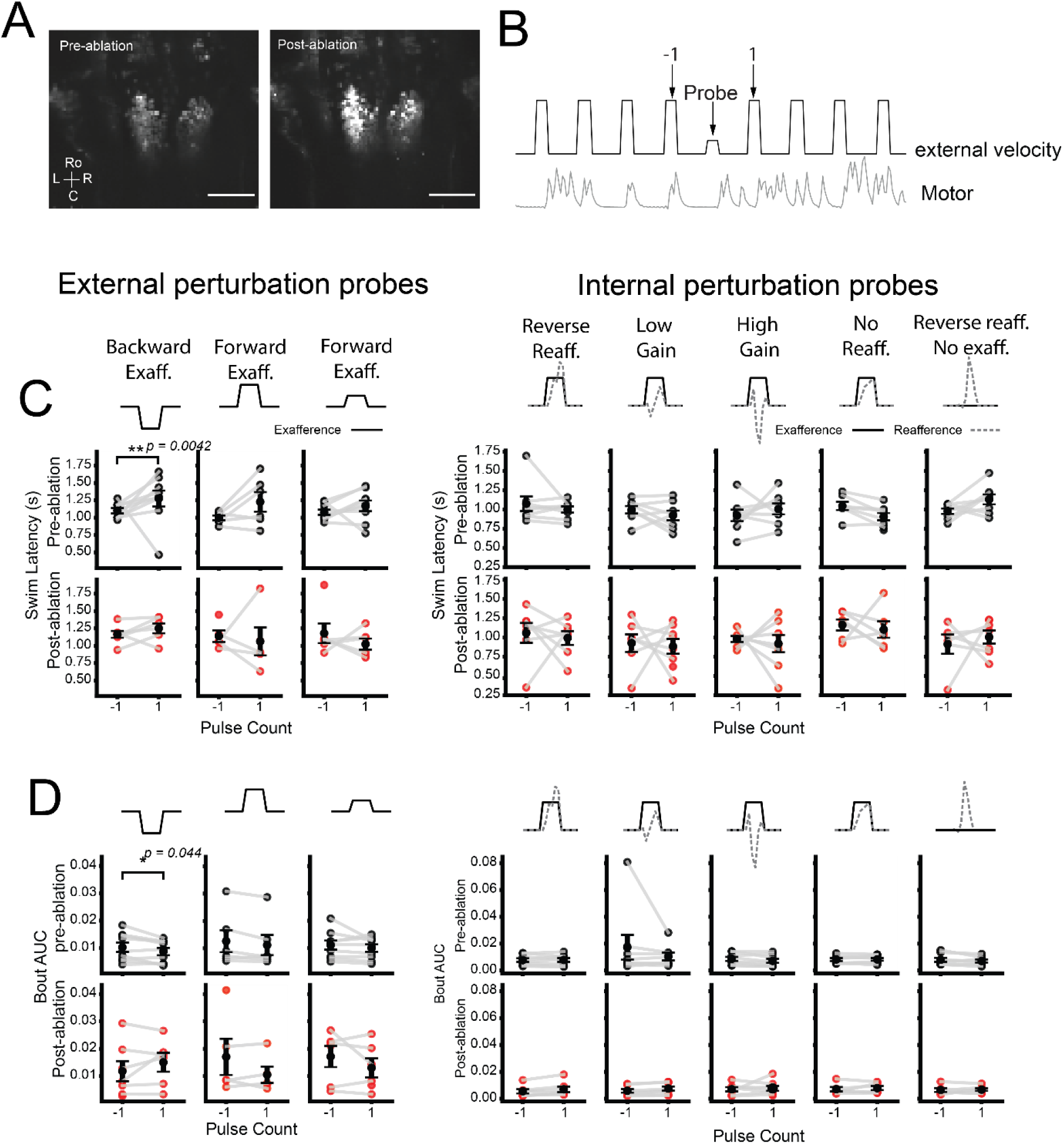
Short-term behavioral responses to external and internal perturbations. **A)** Max projection of the IO taken from 1 fish pre- and post-ablation, top and bottom, respectively. Hyperexcitability and cell death visible in post-ablation. Scale bar 100 µm. B) Top, example trial structure, where the perturbing probe is a change in visual direction, visual velocity, reafference gain or reafference sign inversion. External perturbations all reflect changes in external visual direction or velocity, internal perturbations have consistent external forward velocities, reflecting changes in reafference. C) Left: swim latency for internal perturbation trials. Top rows from pre-ablation trials, bottom rows from post-ablation trials. Right: swim latency for external perturbation trials in the pulse before the probe, -1, and the pulse after probe, 1. D) Left: bout power taken as bout integral for internal perturbation trials. Right: bout power taken as bout integral for external perturbation trials in the pulse before the probe, -1, and the pulse after probe, 1. Top rows from pre-ablation trials, bottom rows from post-ablation trials. Error bars ± SEM.

**Figure S3.**
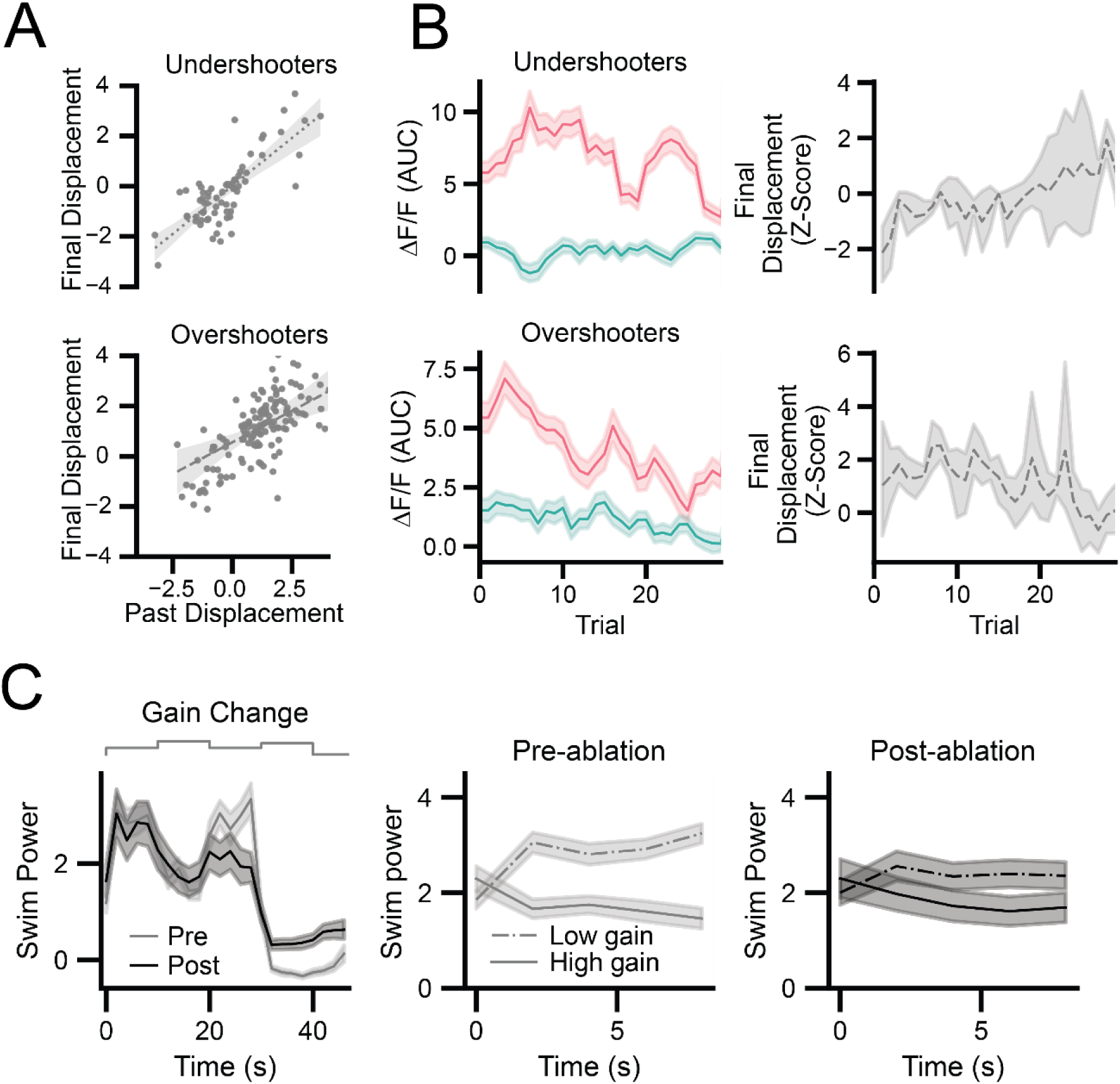
IO responses during adaptation. A) Linear regression of final displacement on current trial compared to previous trial displacement for undershooters (top) and overshooters (bottom). B) Left: Rolling average of forwards and backward- tuned IO neurons over trials for undershooters (top) and overshooters (bottom). Right: Corresponding rolling average of final displacement over trials showing different learning trajectories of undershooters and overshooters. C) The effect of full IO ablation on swim power during the OMR.

**Figure S4.**
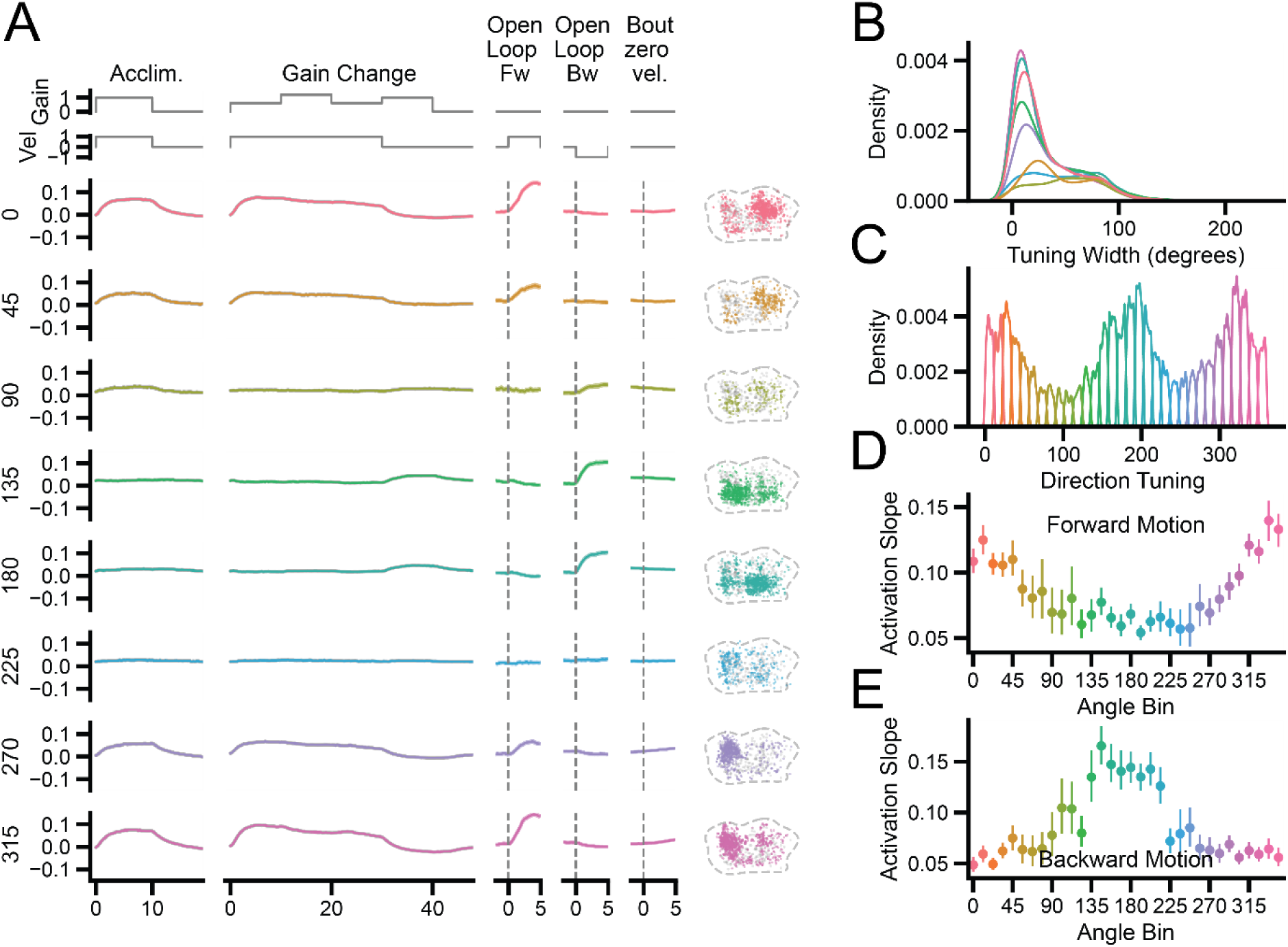
Direction-tuned IO responses during stabilization task. A) IO responses according to direction-tuning bins during the optomotor task, as well as open loop forward and backwards visual motion and bout-triggered events with no visual motion or reafference. B) A kernel-density estimate plot showing the trial-by-trial tuning width distribution. C) A kernel-density plot showing the direction tuning distribution within direction bins (11.25-degree bins). D) The rate of increase in fluorescence (activation slope of ΔF/F) across angle bins during forward visual motion and E) for backward visual motion.

